# Pathways and signatures of mutagenesis at targeted DNA nicks

**DOI:** 10.1101/2021.01.08.425852

**Authors:** Yinbo Zhang, Luther Davis, Nancy Maizels

## Abstract

Nicks are the most frequent form of DNA damage and a potential source of mutagenesis in human cells. By deep sequencing, we have identified factors and pathways that promote and limit mutagenic repair at targeted nicks. BRCA2 inhibits all categories of mutational events at nicks, including indels, SNVs and HDR. DNA2 and RPA promote 5’ resection. Most insertions at nicks consist of a single C incorporated opposite the nick by the translesion polymerase REV1. DNA2 and REV3 inhibit these 1 bp insertions; and DNA2 also inhibits 1 bp deletions. Longer deletions are stimulated by DNA2, REV7 and POLQ. Strikingly, POLQ generates most SNVs at both nicks and double-strand breaks. These results identify mutagenic signatures of DNA2, REV1, REV3, REV7 and POLQ at nicks and highlight the potential for nicks to promote mutagenesis, especially in BRCA-deficient cells.

## Introduction

Nicks are the most frequent form of DNA damage and a potential source of genomic instability in human cells. Nicks can initiate both mutagenesis and homology-directed repair (HDR; reviewed by [1]). Nicks arise naturally in the course of transcription and DNA repair, and they also result from chemical exposures and ionizing radiation (IR), causes 100 nicks for every double-strand break (DSB; [2]). Nicks are significantly less mutagenic than DSBs [3–6]. Nonetheless, nicks have considerable potential to contribute to the overall burden of mutagenesis because of the frequency with which they occur. They may pose a particular threat to genomic stability in tumors treated with IR, which has been a mainstay of cancer therapy for decades, and is currently used to treat over half of solid tumors.

Relatively little is known about pathways that promote mutagenesis at nicks. Evidence derived largely from reporter assays has shown that nicks are normally protected by RAD51, which is loaded onto DNA by BRCA2. Mutagenesis is stimulated in response to reduction in the abundance or activity of RAD51 or BRCA2; of factors that interact with BRCA2 to load RAD51 onto DNA; and by enhancing the activity of the anti-recombinogenic helicase RECQ5, which drives RAD51 off single-stranded DNA [3, 4, 7]. Those same conditions stimulate recombination of nicks with ssDNA donors via an alternative HDR (a-HDR) pathway that uses single-stranded DNA as a donor [3, 4]. These results suggest that, in the absence of protection provided by BRCA2/RAD51, single-stranded regions flanking nicks may become exposed and available to anneal with complementary DNA molecules or to undergo mutagenic repair. However, the factors and pathways that promote mutagenic repair at nicks have not been identified.

To elucidate pathways of mutagenesis at nicks, we have carried out deep sequencing analysis to characterize mutagenic events at targeted nicks in human cells. Here we show that deletions and SNVs are asymmetrically distributed around a nick site, an asymmetry that may reflect the activity of DNA2/RPA, which promote 5’ resection at nicks. DNA2 also contributes to long deletions, but inhibits both 1 bp insertions and 1 bp deletions at nicks. The majority of insertions at nicks are of a single C on the strand opposite the nick, a signature that was greatly reduced in REV1-depleted cells, and thus reflects the dCMP transferase activity of REV1 translesion polymerase. REV3 inhibited these and other insertions; and REV7 inhibited deletions. POLQ plays an unanticipated role in promoting SNVs at nicks, and also at DSBs. These results define pathways of mutagenesis active at nicks and, and define mutagenic signatures of factors that carry out or regulate nick repair. They also highlight the potential for nicks to promote mutagenesis, especially in BRCA-deficient cells.

## Results

### BRCA2 depletion increases the frequency of mutations at nicks (but not DSBs)

We examined frequencies of mutagenic events, including deletions, insertions and SNVs, at nicks and DSBs targeted to a site in the non-transcribed strand of the endogenous CD44 gene in human U2OS cells (**Fig 1A**). U2OS is a human osteosarcoma line with wild-type P53 that is frequently used to study DNA repair. CD44 encodes a cell-surface glycoprotein that is expressed in many cell types but not required for proliferation in cell culture. Surface CD44 is readily detected by antibody staining, making it possible to verify that observations document events at a transcribed gene by using flow cytometry to assay gene expression. DNA nicks and DSBs were targeted to exon 1 of the CD44 gene by Cas9D10A or Cas9, respectively, and CRISPR gRNA 4. Genomic DNA was isolated from 10,000-30,000 cells and PCR-amplified, and deletions, insertions and SNVs were scored within a 65 bp window by amplicon sequencing on an Illumina platform. To establish the combined error frequencies of PCR amplification, sequencing, and computational analysis, control experiments analyzed uncleaved DNA. Indels and SNVs were assessed using a computational strategy that greatly reduced PCR and sequencing errors by making a consensus call for groups of DNA molecules that contained the same unique molecular identifiers (UMI; see Methods). Absolute frequencies of indels and SNVs varied among experiments, but the effects of depletion of specific factors were reproducible. Results are presented for representative sequenced libraries.

**Fig 1.**
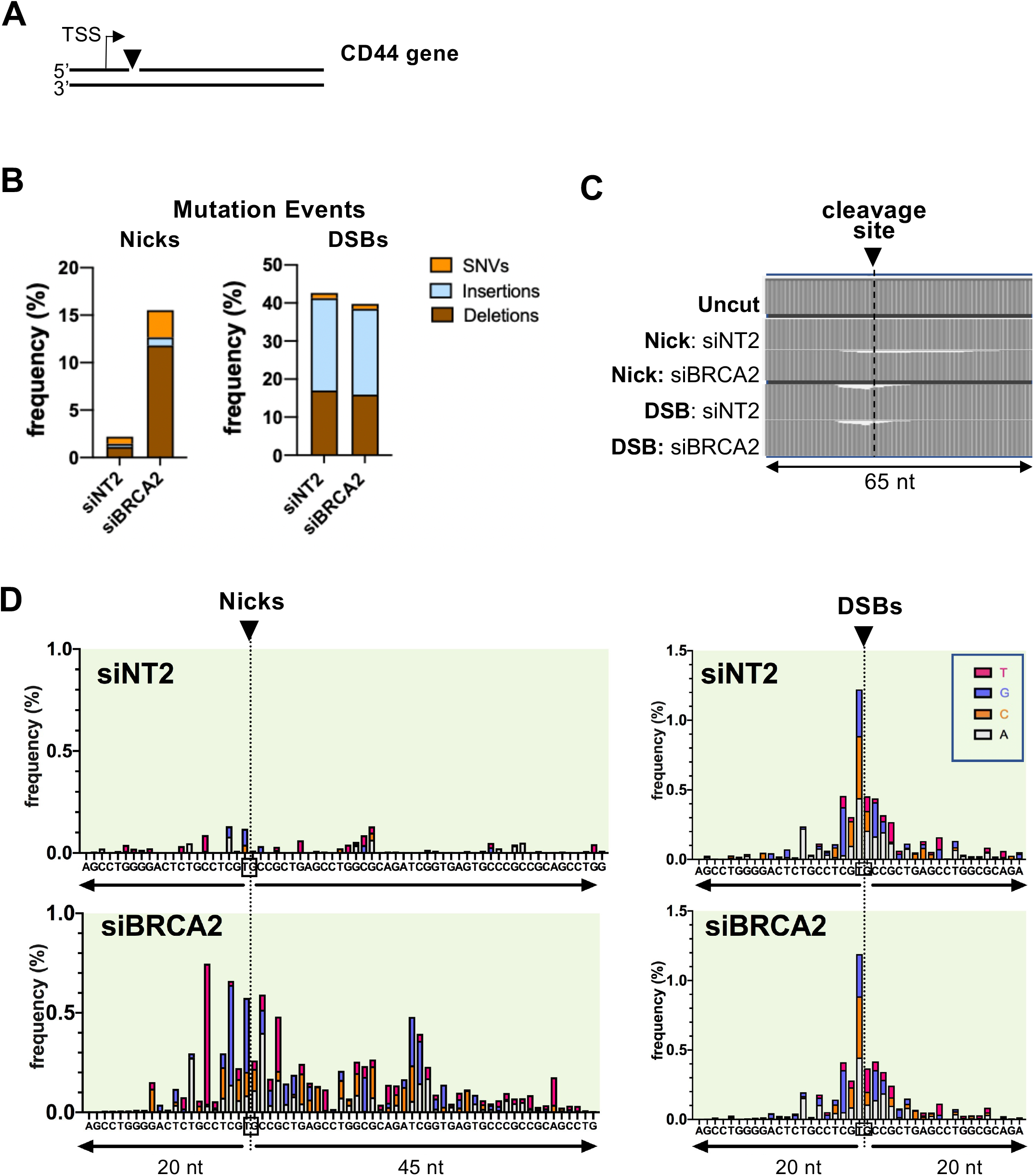
Depletion of BRCA2 increases the frequency of mutations at nicks. **(A)** Diagram of site targeted for cleavage in CD44 gene of human U2OS cells, showing transcription start site (TSS) and the site on the non-transcribed strand targeted for cleavage by gRNA 4 (arrowhead). **(B)** Frequencies of mutation events at nicks and DSBs targeted by gRNA4, as determined by amplicon sequencing. Deletions, insertions and single nucleotide variations (SNVs) were scored in control cells (siNT2) or cells depleted for BRCA2. **(C)** Maps of the fractional decrease in the number of bases at each position within the indicated 65 nt region spanning the gRNA4 cleavage site in U2OS cells. DNA was uncut, or targeted for nicks or DSBs, in control cells (siNT2) or cells depleted for BRCA2, as indicated. **(D)** Maps of positions and spectra of SNVs within the indicated regions spanning nicks or DSBs in U2OS cells treated with control siNT2 or siBRCA2. Arrowheads and dashed lines below them indicate the site of the nick or DSB, which targets the DNA backbone between the nucleotides boxed in the sequence below.

At nicks, the frequency of mutagenic events was considerably lower than at DSBs (2.2% vs. 42%; 19-fold; **Fig 1B**). At nicks, most mutagenic events were deletions, with fewer SNVs and few insertions; while at DSBs most were insertions, with somewhat fewer deletions and relatively few SNVs. Depletion of BRCA2 increased the frequency of mutagenic events at nicks 7-fold (p <0.01), but had no significant effect at DSBs (p=0.985).

Maps of the fractional decrease in the number of base calls at each position showed that, in cells depleted of BRCA2, deletions exhibited an asymmetric distribution with respect to the nick site, extending from 10 bp 5’ (upstream) to 30 bp 3’ (downstream); while at DSBs, deletions were clustered within approximately 10 bp on either side of the cleavage site (**Fig 1C**). SNVs also exhibited asymmetric distribution around the target site of nicks, but not of DSBs (**Fig 1D**). This asymmetry was especially clear in the map of SNVs in BRCA2-depleted cells, where SNVs extended from 20 bp 5’ (upstream) to 45 bp 3’ (downstream) of the nick (**Fig 1D** left). In contrast, SNVs were clustered near the target site of DSBs and extended no more than 10-15 bp to either side), in a pattern relatively unaffected by BRCA2 depletion (**Fig 1D** right).

### DNA2 and RPA1 promote resection resulting in a-HDR or mutagenesis at nicks

The asymmetric distribution of deletions and SNVs around nicks suggested that 5’-resection may be an early step in mutagenesis. We tested this by using the Traffic Light (TL) reporter [8] to determine the effect of depletion of candidate nucleases on frequencies of a-HDR at nicks in cells provided with a single-stranded oligonucleotide donor and co-depleted for BRCA2. Frequencies of canonical (c-HDR) at DSBs in cells provided with a dsDNA plasmid donor were quantified in parallel. Frequencies of a-HDR at nicks were reduced in response to depletion of DNA2, but unaffected by depletion of EXO1 or MRE11; while frequencies of c-HDR at DSBs targeted to the same site were reduced in response to depletion of MRE11, DNA2, and to a lesser extent EXO1 (**S1A Fig**). A similar 2-fold reduction in a-HDR frequencies was evident at nicks targeted to either the transcribed or non-transcribed strand of the reporter construct and supported by ssDNA donors complementary to either the nicked (cN) or intact (cI) strand (**S1B Fig**). DNA2 has both nuclease and motor activities which enable it to expose and resect 5’ ends and flaps in DNA replication and recombination [9, 10]. This 5’ resection activity is consistent with the asymmetric distribution of deletions and mutations around the site of a nick (**Fig 1D**). It is also consistent with the involvement of DNA2 in a-HDR at nicks supported by a ssDNA donor complementary to either the nicked or intact strand (as diagrammed in **S1C Fig**).

The trimeric factor RPA coats single-stranded gaps and free DNA ends exposed by DNA2 to prevent annealing and structure formation and coordinate end resection [11–13]. However, while cell cycle was unperturbed by depletion of DNA2, depletion of RPA1, the largest subunit of RPA, resulted in cell cycle arrest (**S2A Fig)**. We therefore pursued an alternative strategy. The *S. cerevisiae* mutant Rfa1-t11 (K45E) supports normal replication but causes a defect in both mitotic and meiotic recombination [14–16]. Human RPA1 derivatives with mutations mapping near *S. cerevisiae* K45E were previously shown to support replication but were not analyzed for function in recombination [17, 18]. By alignment we identified human RPA1 residues R41 or R43 as likely counterparts of *S. cerevisiae* K45 (**Fig 2A**). Based on this, we generated human RPA1 expression clones bearing the R41E and R43E single mutations as well as an R41/43E double mutation, and tested their effects on replication and HDR. Ectopic expression of RPA1-R41E, RPA1-R43E, or RPA1-R41/43E did not affect cell cycle (**S2B Fig**), but inhibited both a-HDR at nicks and c-HDR at DSBs (**Fig 2B**). Expression of either RPA1-R43E or RPA1-R41/43E reduced both a-HDR and c-HDR more than 5-fold. Note that inhibition was evident in cells expressing endogenous RPA, so these mutants exert a dominant negative effect on recombination. The RPA1-R43E single mutant was used for further analysis.

**Fig 2.**
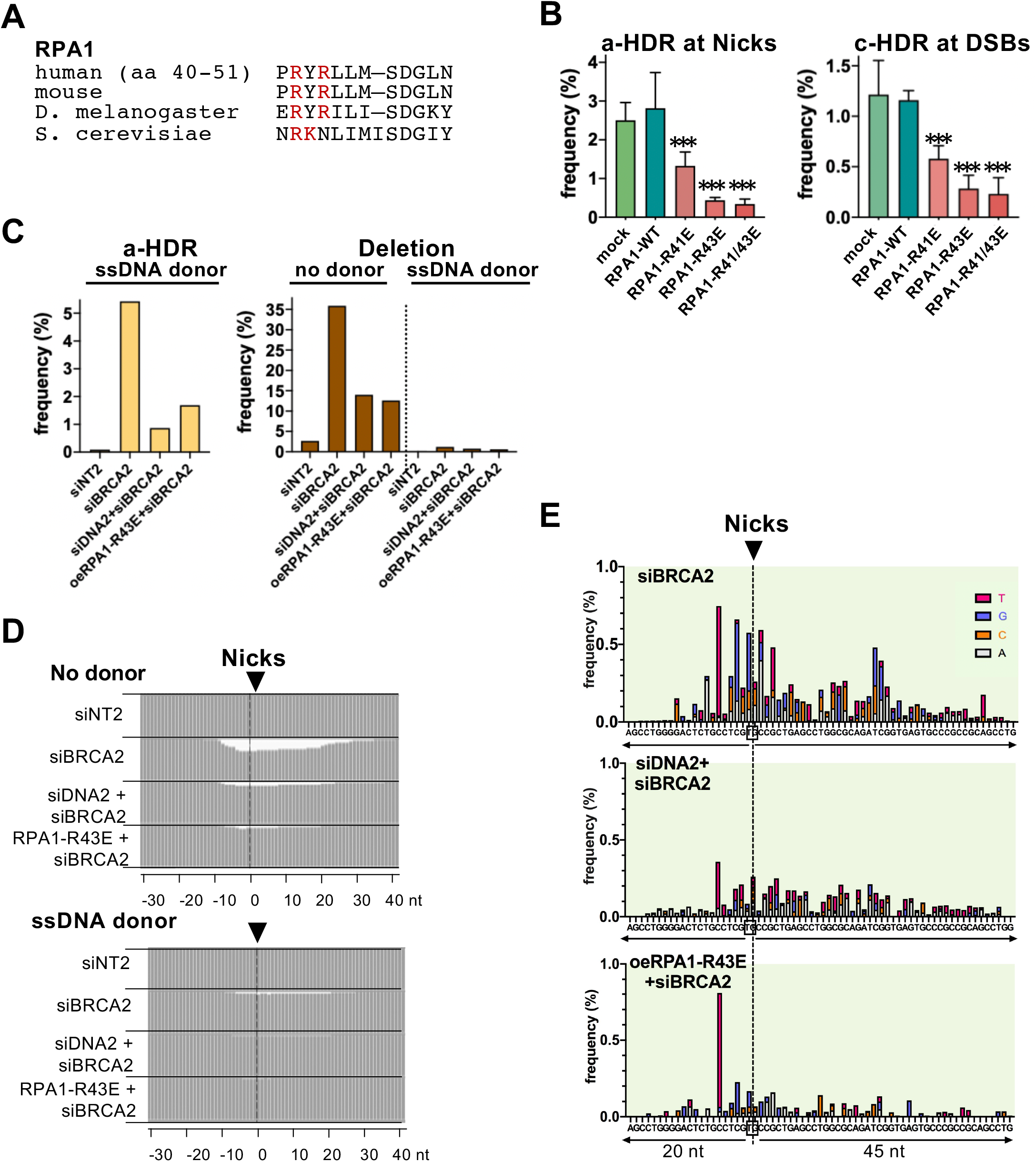
DNA2 and RPA1 promote resection resulting in a-HDR or mutagenesis at nicks. **(A)** Alignment of RPA1 protein sequences in the conserved N-terminal region to which the *S. cerevisiae* Rfa1-t11 (K45E) mutation [14] maps. **(B)** Frequencies of a-HDR at nicks or c-HDR at DSBs as determined by reporter assays in 293 TL cells ectopically expressing RPA1 WT or its mutant derivatives. Frequency values represent the mean ± SEM from a representative experiment; *** indicates p<0.001 for the frequency difference between the indicated sample and sample expressing RPA1-WT. **(C)** Frequencies of a-HDR at nicks targeted to the CD44 gene in U2OS cells ectopically expressing RPA1-WT or its mutant derivatives and provided with a ssDNA donor for a-HDR by sequence analysis. **(D)** Maps of the fractional decrease in the number of base calls at each position within the indicated region spanning the gRNA4 cleavage site in U2OS cells in cells treated as indicated and lacking or provided with a ssDNA donor complementary to the nicked strand (cN) to support a-HDR. **(E)** Maps of positions and spectra of SNVs within the indicated 65 bp region spanning nicks targeted by gRNA4 to the CD44 gene in U2OS cells treated as indicated.

The roles of DNA2 and RPA in mutagenesis and a-HDR were then analyzed by amplicon sequencing. Sequence analysis showed that depletion of BRCA2 greatly stimulated a-HDR frequencies, which were reduced by co-depletion of DNA2 or by ectopic expression of RPA1-R43E (6.2- and 3.2-fold, respectively; **Fig 2C** left). Deletion frequencies were high in BRCA2-depleted cells in the absence an HDR donor, and reduced in BRCA2-depleted cells by co-depletion of DNA2 or by ectopic expression of RPA1-R43E (2.6- and 2.0-fold, respectively); and provision of an HDR donor reduced deletion frequencies more than 20-fold (**Fig 2C** right). Maps of the fractional decrease in the number of bases at each position showed that the extent of deletions 3’ (downstream) of the nick site was reduced in response to either siDNA2 treatment or ectopic expression of RPA1-R43E (**Fig 2D**). In cells depleted of BRCA2, co-depletion of DNA2 or ectopic expression of RPA1-R43E reduced the frequencies of SNVs at nicks to 60% and 38%, relative to depletion of BRCA2 alone (**Fig 2E**). These results support the view that availability of nicks as substrates for HDR or mutagenesis depends upon both DNA2 and RPA, and further suggest that pathways of mutagenesis and a-HDR compete for the same DNA substrate.

### Cell cycle regulates DNA2/RPA-mediated resection at nicks

DNA2 and RPA are regulated by cell cycle and most active in S phase [10]. c-HDR at DSBs, which depends on both these factors, is most efficient in S phase [19, 20], and the dependence of a-HDR at nicks on these factors (**Fig 2**) suggested that a-HDR at nicks might also be most efficient in S phase. To test this, we assayed a-HDR and c-HDR in cells in which the nuclear activities of Cas9D10A, Cas9 or RPA were restricted to G1 or S/G2 phase of cell cycle. Expression constructs were generated bearing these factors fused to degrons derived from the CDT1 or geminim (GEM) cell cycle regulators [21], and the predicted restriction to G1 or G2/S phase, respectively, was confirmed by flow cytometry (**S3 Fig**). Relative efficiencies of HDR initiated during G1 or S phase were then quantified using the TL reporter assay (**Fig 3**). G1 phase nicks initiated a-HDR more efficiently than S/G2 phase nicks (generated by Cas9D10A-CDT1 and Cas9D10A-GEM, respectively; **Fig 3A** left). In contrast, as previously reported [19, 20], S/G2 phase DSBs initiated c-HDR more efficiently than G1 phase DSBs (generated by Cas9-GEM and Cas9-CDT1, respectively; **Fig 3A** right).

**Fig 3.**
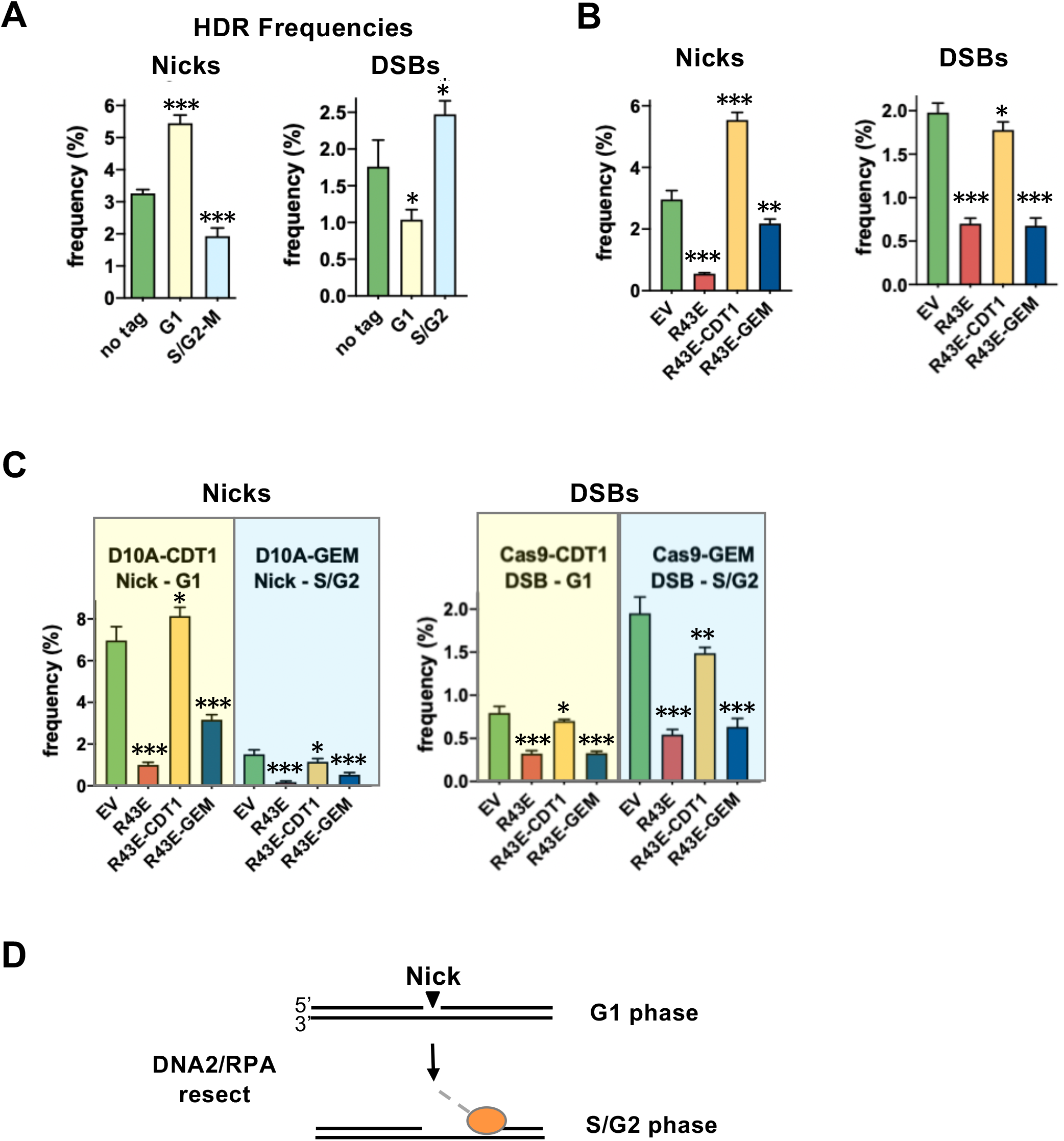
Cell cycle regulates HDR at nicks. **(A)** Effect on HDR frequencies of restriction of nicks or DSBs to G1 or S/G2 phase. HDR frequencies were determined using the TL reporter in 293T TL cells transfected with untagged Cas9D10A or Cas9, or with derivatives bearing CDT1 or GEM tags to restrict cleavage to G1 or S/G2 phase, respectively. Cleavage was targeted by gRNA9 and supported by a cN ssDNA for a-HDR (nicks) or a plasmid donor for c-HDR (DSBs). Frequency values represent the mean ± SEM from a representative experiment; *, **, and *** indicate p<0.05, p<0.01, and p<0.001, respectively, for the frequency difference between indicated sample and sample transfected with untagged construct. **(B)** Effect on HDR frequencies at nicks and DSBs of restriction of the inhibitory activity of RPA1-R43E to G1 or S/G2 phase. Frequency values represent the mean ± SEM from a representative experiment; *, **, and *** indicate p<0.05, p<0.01, and p<0.001, respectively, for the frequency difference between indicated sample and sample transfected with empty vector (EV). **(C)** Effect on HDR frequencies at nicks and DSBs of cell cycle restriction of both cleavage and RPA1-R43E activities. Results obtained when cleavage was restricted to G1 or S/G2 distinguished by yellow or blue shading. Other details as in panel B. **(D)** Diagram of how a nick generated in G1 phase may persist to undergo resection later in cell cycle. While the analysis reported here does not directly connect the importance of RPA activity in S/G2 phase to its well-studied role in DNA2/RPA-mediated resection, it is consistent with that function.

Cell cycle dependence of RPA function was queried by assessing HDR in cells in which the dominant negative RPA1-R43E mutant was expressed bearing either a CDT1 or GEM tag. RPA1-R43E-CDT1 will inhibit HDR in G1 phase, and RPA1-R43E-GEM will inhibit HDR in S/G2 phase, so if HDR occurs preferentially in S/G2 phase, HDR frequencies are predicted to be reduced in cells expressing RPA1-R43E-GEM relative to control cells. This pattern was evident not only at DSBs (**Fig 3B** right), as predicted by previous results [19, 20], but also at nicks (**Fig 3B** left). Thus, RPA promotes HDR at both nicks and DSBs more effectively in S/G2 phase than in G1 phase.

At nicks, a-HDR frequencies were unexpectedly higher in cells expressing RPA-R43E-CDT1 than in control cells, an effect not evident at DSBs (**Fig 3B**). A possible explanation for this is that nicks generated in G1 phase undergo HDR most efficiently if they persist into S/G2 phase for repair. To address this possibility, HDR was assayed at nicks and DSBs initiated in either G1 or S/G2, by Cas9D10A or Cas9 bearing CDT1 or GEM degron tags, in cells expressing dominant negative RPA1-R43E or its derivatives bearing CDT1 or GEM degron tags (**Fig 3C**). In all cases, HDR frequencies were higher in cells expressing RPA1-R43E-CDT1, which inhibits HDR in G1 phase, than in cells expressing RPA1-R43E-GEM, which inhibits HDR in S/G2 phase. Thus G1 phase nicks may persist to undergo repair later in cell cycle.

### DNA2 inhibits 1 bp deletions and insertions at nicks, but promotes longer deletions

Amplicon sequencing was used to examine the frequencies and lengths of deletions and insertions at nicks and DSBs in cells depleted for BRCA2 and DNA2 (**Fig 4**). In control cells (siNT2), the frequency of deletions was 20-fold lower at nicks than at DSBs (**Fig 4A**). At nicks, depletion of BRCA2 stimulated deletion frequencies more than 10-fold, and most deletions were >20 bp in length; while at DSBs, depletion of BRCA2 had little effect on deletion frequencies, and most deletions were <10 bp. At nicks, co-depletion of DNA2+BRCA2 reduced the overall deletion frequency nearly 2-fold relative to depletion of BRCA2 alone, reflecting reduced frequencies of longer deletions (**Fig 4A** left). At DSBs, the overall deletion frequency was also reduced nearly 2-fold by depletion of DNA2, but due primarily to a reduction in short deletions (**Fig 4A** right). Separate analysis of deletions of 1-6 bp in length revealed that at nicks, co-depletion of DNA2+BRCA2 caused a 6-fold ***increase*** in the frequency of 1 bp deletions, but reduced the frequency of deletions of 2-6 bp; while at DSBs, DNA2 depletion reduced the frequencies of all 1-6 bp deletions (**Fig 4B**).

**Fig 4.**
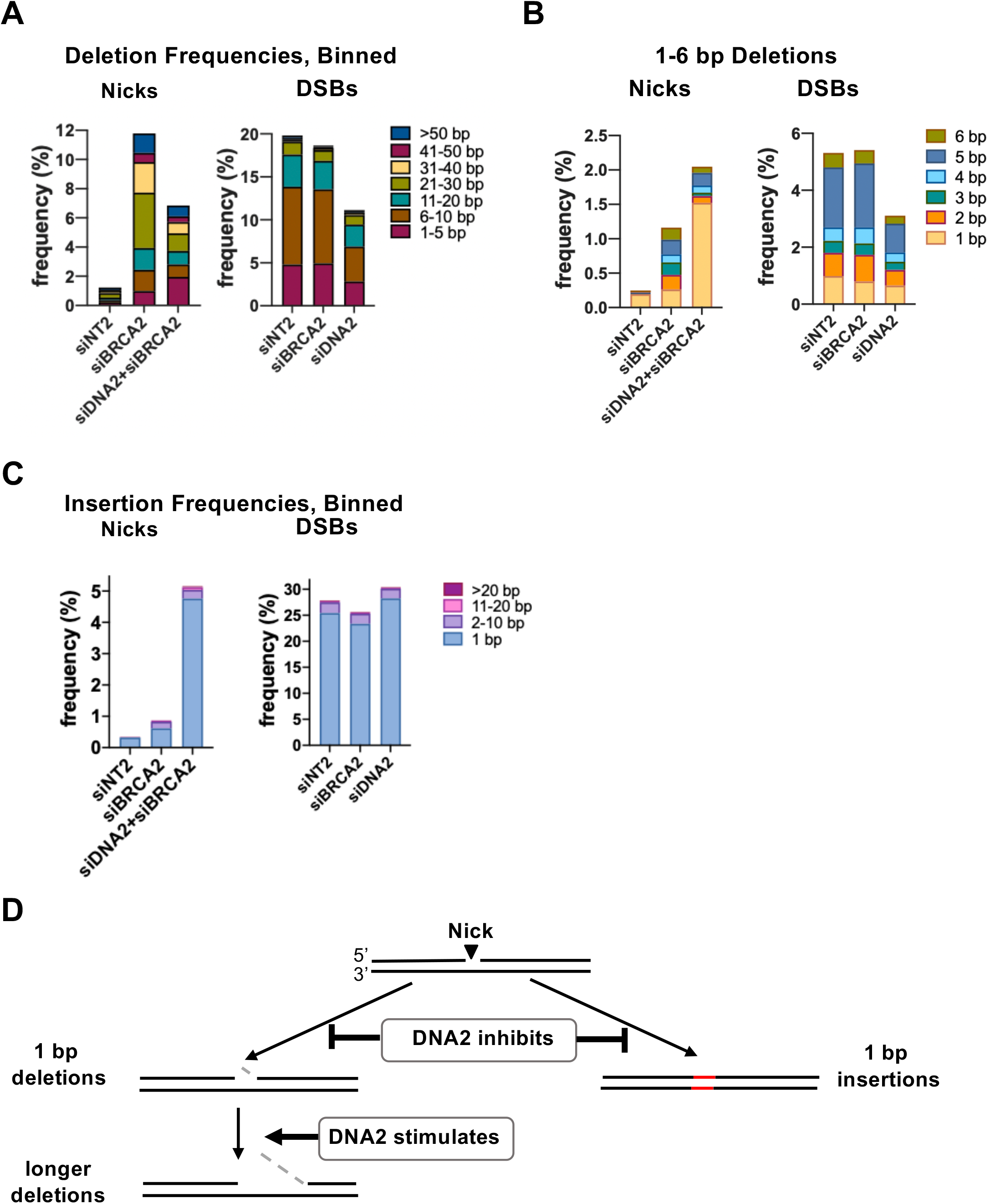
DNA2 inhibits 1 bp indels but promotes longer deletions at nicks. **(A)** Effects of depletion of BRCA2 and/or DNA2 on frequencies of deletions of indicated lengths at nicks or DSBs targeted to the CD44 gene in U2OS cells, determined by amplicon sequencing. **(B)** Effects of depletion of BRCA2 and/or DNA2 on frequencies of 1-6 bp deletions at nicks or DSBs targeted to the CD44 gene in U2OS cells. **(C)** Effects of depletion of BRCA2 and/or DNA2 on frequencies insertions of indicated lengths at nicks or DSBs targeted to the CD44 gene in U2OS cells. **(D)** Diagram of regulation of deletions and insertions by DNA2. Resected DNA is indicated by gray dashes.

Insertions were rare (<1%) at nicks in control cells (siNT2), and far more frequent at DSBs (**Fig 4C**). At nicks, insertion frequencies increased 3-fold in response to depletion of BRCA2 alone, and 17-fold in response to co-depletion of DNA2+BRCA2, reaching 5%; while at DSBs, depletion of BRCA2 or DNA had little effect on insertion frequencies. Surprisingly, the great majority of insertions at both nicks and DSBs were 1 bp in length, in all conditions tested.

Taken together, the results above identify an unexpectedly complex role for DNA2 at nicks (**Fig 4D**). DNA2 promotes deletions at nicks, as it does at DSBs; however, at nicks this effect is restricted to deletions longer than 1 bp, and DNA2 specifically inhibits 1 bp deletions. DNA2 also inhibits the 1 bp insertions that constitute the majority of insertion events at nicks. At DSBs, 1 bp insertions frequently result from fill-in reactions at 5’ overhangs generated as a result of staggered cleavage by Cas9 [22–24]. However, there are no overhangs to fill in at a nick, so some other mechanism must generate these 1 bp insertions. This mechanism is identified below.

### REV1 promotes 1 bp insertions at nicks, and REV3 inhibits insertions

To identify pathways that contribute to mutagenic repair, amplicon sequencing was used to analyze libraries of sequences at nicks in U2OS cells depleted for four factors, REV1, REV3, REV7 or POLQ, along with BRCA2. The roles of each of these factors has previously been extensively studied. REV1 is a Y family polymerase that functions as a dCMP transferase, inserting a single C to bridge unusual structures during translesion synthesis [25]. REV3 and REV7 (also known as MAD2L2) are two components of the four-subunit B-family DNA polymerase POLζ (REV3/REV7/POLD2/POLD3), and they are recruited by REV1 and exchange with it to extend a distorted DNA primer-template [26]. REV7 also functions independently to protect telomeres and DSBs from resection and to regulate NHEJ at DSBs as a component of the shieldin complex [27–33]. POLQ encodes POLθ, a DNA helicase and translesion polymerase that promotes repair of DSBs formed by a variety of mechanisms, including replication fork stalling, targeted DNA cleavage and ionizing radiation, by binding to and extending annealed short (2-6 bp) duplex regions; maintains genomic stability by preventing interhomolog recombination, which can result in loss-of-heterozygosity; and, conversely, promotes insertions and random chromosomal integration [34–40]. Libraries were generated in parallel from identically depleted cells targeted with nicks or DSBs, to facilitate comparison with previously described activities and to control for depletion.

At nicks (**Fig 5A** left), insertion frequencies increased several-fold in response to depletion of BRCA2. Co-depletion of REV1+BRCA2 caused a 3-fold reduction in the frequency of all insertions relative to BRCA2-depleted cells (0.29% vs 0.87% respectively) and eliminated 1 bp insertions. This immediately implicated REV1 as the source of 1 bp insertions. Insertion frequencies increased nearly 3-fold in response to depletion of REV3+BRCA2, relative to depletion of BRCA2 alone (2.44% vs 0.87%, respectively), suggesting that REV3 normally inhibits 1 bp insertions. Co-depletion of REV7+BRCA2 had little effect on insertion frequencies. Co-depletion of POLQ+BRCA2 caused a modest increase in the frequency of 1 bp insertions, and a modest reduction in longer insertions.

**Fig 5.**
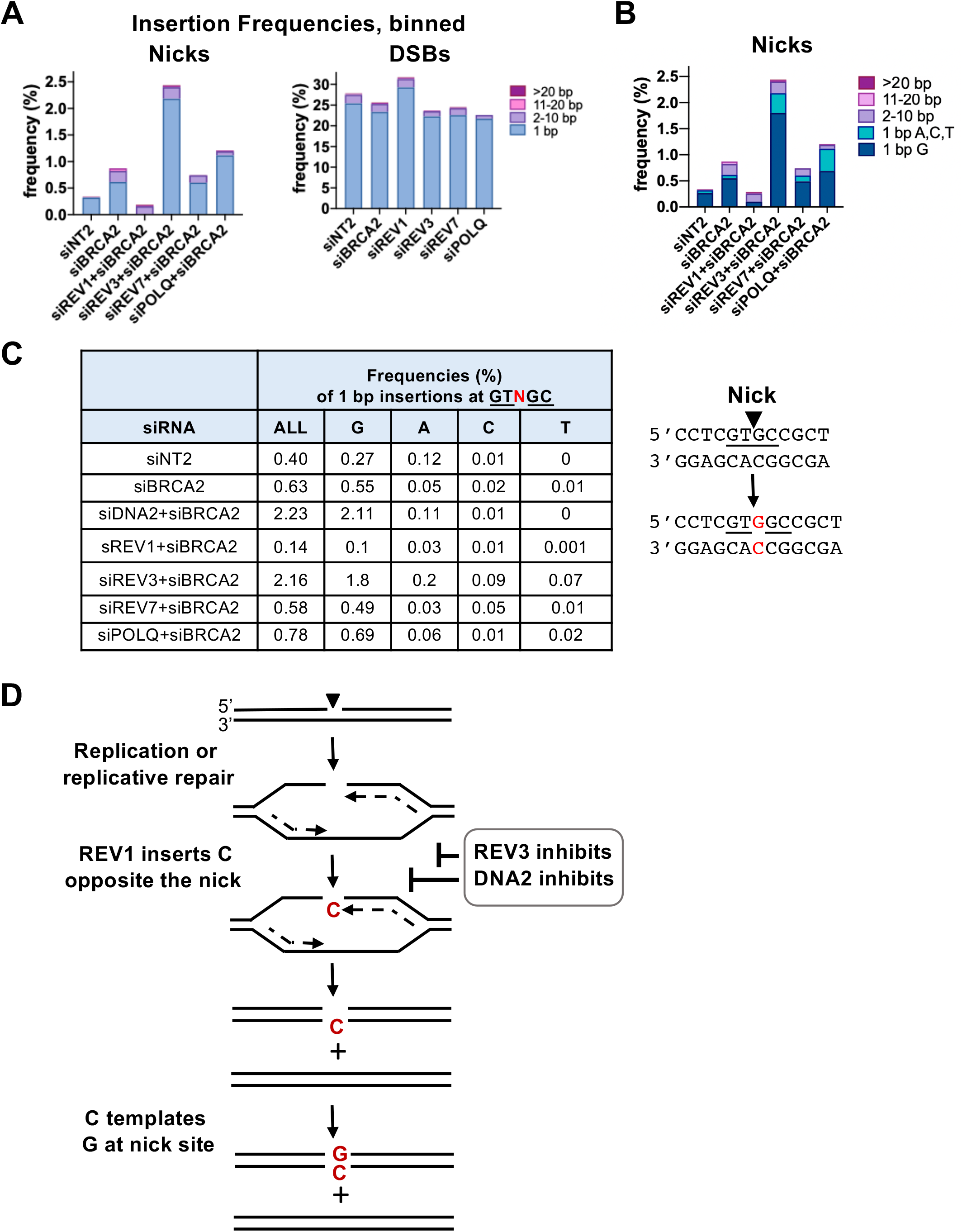
REV1 promotes and REV3 and DNA2 inhibit 1 bp insertions at nicks. **(A)** Effects of depletion of indicated factors on frequencies of insertions of indicated lengths at targeted nicks and DSBs at the CD44 gene in U2OS cells, as determined by amplicon sequencing. **(B)** Effects of depletion of indicated factors on frequency of molecules bearing a single bp insertion of A,C,T or G. Sequences are shown for the nicked strand **(C)** Left, frequencies of 1 bp insertions of all nucleotides and G, A,T or C at the nick site. Right, diagram of the majority of sequence changes due to 1 bp insertion; nick site, underlined, insertion, red font. **(D)** Diagram of how REV1 may insert C opposite a nick during replication, which then templates a G on the opposite strand, resulting in insertion of a G at the nick. Inhibition by REV3 and DNA2 indicated by lines, with relative lengths reflecting relative magnitudes of inhibition. Replicating DNA indicated by dashed black arrows; insertion in red font.

Insertion frequencies at DSBs (**Fig 5A** right) were relatively unaffected by depletion of BRCA2, and increased modestly upon depletion of REV1. At DSBs, depletion of either REV3 or REV7 caused a reduction in insertion frequencies of similar magnitude (15%), which may reflect function of these factors in concert as components of POLζ. This contrasts with the striking increase in insertion frequencies at nicks in REV3-but not REV7-depleted cells (**Fig 5A** left), and suggests that REV3 and REV7 function independently at nicks, although not at DSBs. At DSBs, depletion of POLQ caused a modest increase in 1 bp insertions and a decrease in longer insertions.

The results in **Fig 5A** suggest that REV1 is responsible for most or even all 1 bp insertions at nicks. REV1 is a translesion polymerase with dCMP transferase activity, making it possible to directly test this hypothesized role. The sequencing output “reads” the strand targeted by the nick, so an insertion of C on the nicked strand it will appear as a C in the sequence data file, and an insertion in the opposite strand will appear as a G. As shown in **Fig 5B**, almost all insertion events could be accounted for by insertion of a single G in the nicked strand, in control (siNT2) cells and in all other samples tested, except cells depleted for REV1. These insertions mapped to the site of the nick, and not nearby positions (**Fig 5C** left; **S4 Fig**), changing the duplex DNA sequence as shown (**Fig 5C** right). These results establish that REV1 generates 1 bp insertions at nicks, and give rise to a model in which REV1 inserts a single C opposite the site of the nick, which then templates addition of a G on the opposite strand (**Fig 5D**).

Sequence analysis further identified unexpected roles for REV3 and DNA2 in suppressing 1 bp insertions. Co-depletion of either REV3+BRCA2 (**Fig 5A,C**) or DNA2+BRCA2 (**Fig 4C**, **5C**) increased the overall insertion frequency and the frequency of insertions of G at the site of the nick on the nicked strand. The evidence that DNA2 inhibited insertions at nicks while stimulating long deletions (**Fig 4**) raises the possibility that resection at the nick by DNA2 generates a gap that becomes the source of deletions, rather than a substrate for REV1-mediated insertion.

### REV7 and POLQ promote deletions longer than 1 bp at nicks

Analysis of deletion frequencies by amplicon sequencing showed that, at nicks, there was no effect of co-depletion of REV1 or REV3 and BRCA2 relative to depletion of BRCA2 alone, while co-depletion of REV7 or POLQ and BRCA2 reduced deletion frequencies (31% and 47%, respectively; **Fig 6A** left). At DSBs, reductions in deletions frequencies in the range of 10%-40% resulted from depletion of REV1, REV3, REV7 or POLQ (**Fig 6A** right).

**Fig 6.**
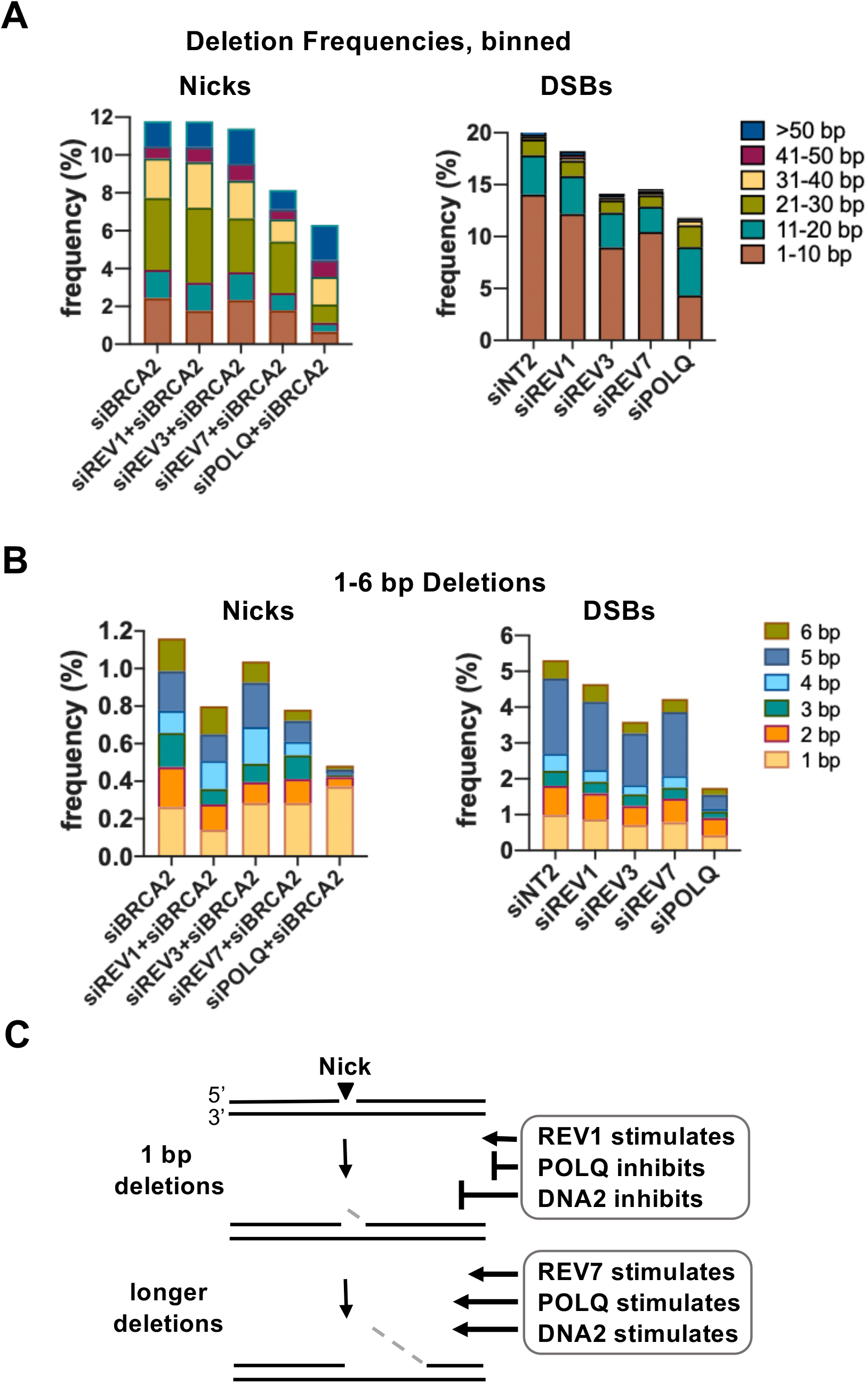
REV7 and POLQ promote deletions at nicks. **(A)** Effects of depletion of indicated factors on frequencies of deletions, binned by tens, at targeted nicks and DSBs in the CD44 gene of U2Os cells. **(B)** Effects of depletion of indicated factors on frequencies of 1-6 bp deletions. **(C)** Diagram of distinct regulation of 1 bp and longer deletions by REV1, REV7, POLQ and DNA2. Stimulation and inhibition indicated by lines, with relative lengths reflecting relative magnitudes of stimulation or inhibition.

Separate examination of short deletions (1-6 bp, **Fig 6B**) showed that at nicks, co-depletion of BRCA2 and REV1, REV7 or POLQ reduced the overall frequencies of deletions in this range, but with contrasting effects on 1 bp deletions, which were reduced 2-fold by REV1 depletion, unaffected by REV7 depletion, and increased nearly 2-fold by POLQ depletion. Depletion of these same factors, most notably POLQ, also reduced the frequencies of 1-6 bp deletions at DSBs. These results and the analysis of the effects of co-depletion of BRCA2 and DNA2 at nicks (**Fig 4B,D**) both suggest that 1 bp deletions arise or are processed differently from longer deletions at nicks: 1 bp deletions are stimulated by REV1 and inhibited by POLQ and especially DNA2, and longer deletions are stimulated by REV7 and especially POLQ and DNA2 (**Fig 6C**).

### POLQ contributes to most SNVs at nicks

SNV frequencies at nicks and DSBs were determined as described in Methods and normalized to siBRCA2 or siNT2-treated cells, respectively, to facilitate comparisons of the effects of each factor (**Fig 7A**). At nicks, frequencies of SNVs increased 4-fold in response to depletion of BRCA2, accompanied by a switch to a predominance of transversion mutations (from 25% to 70%). Co-depletion of REV1, REV3 or REV7 along with BRCA2 reduced SNV frequencies 15-30% relative to cells depleted of BRCA2 alone, but had relatively little effect on the fraction of transversion mutations. At DSBs, depletion of BRCA2 had little effect on SNV frequency or the fraction of transversions relative to control (siNT2) cells, and depletion of REV1, REV3 or REV7 reduced SNV frequencies with little effect on the fraction of transversions. Strikingly, SNV frequencies at both nicks and DSBs were strongly reduced in response to depletion of POLQ, and the fraction of transversions diminished at nicks, though not at DSBs.

**Fig 7.**
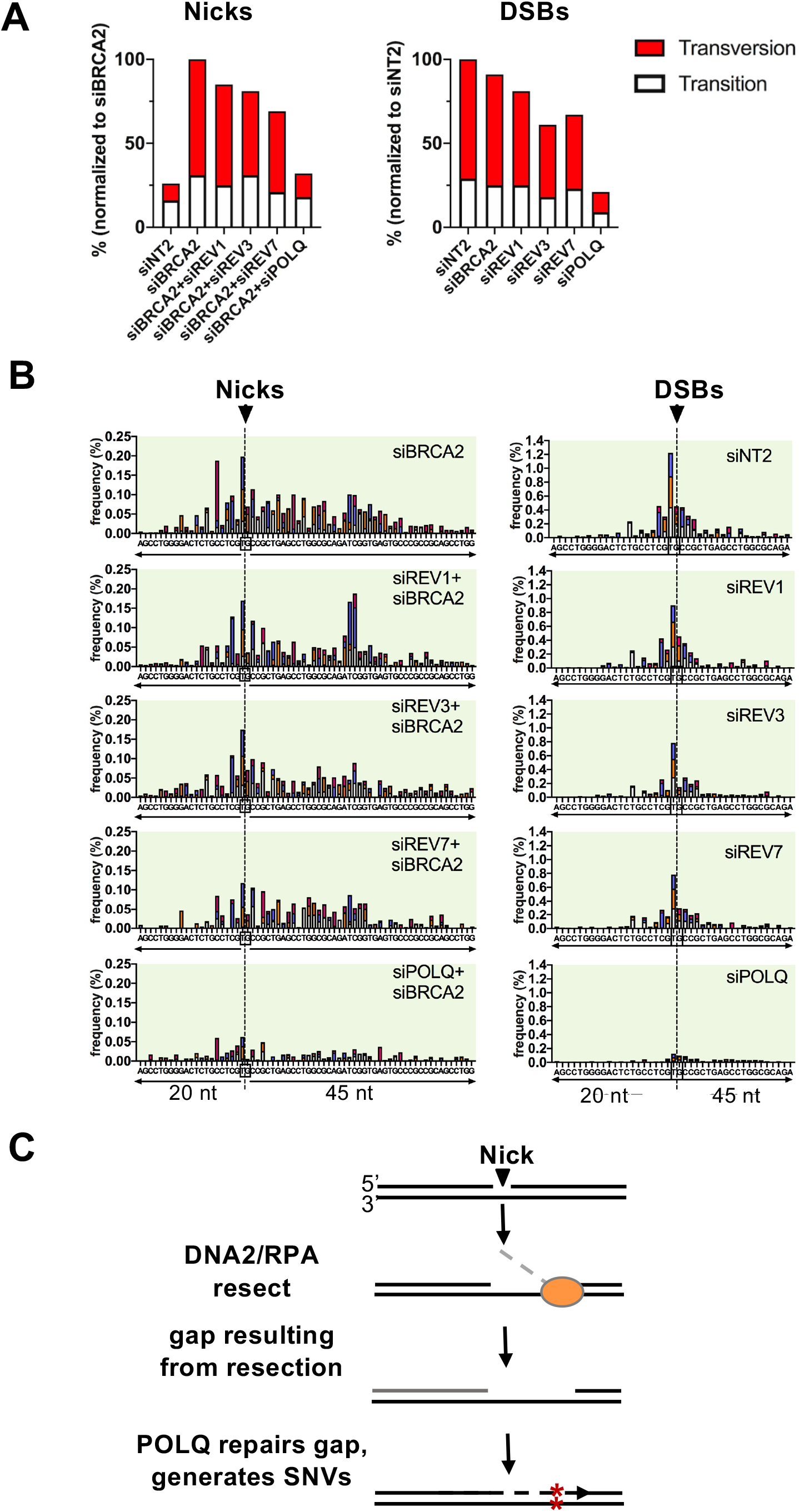
SNVs. **(A)** Frequencies of SNVs at nicks and DSBs in the CD44 gene of U2OS cells depleted for indicated factors. To facilitate comparison of effects of different factors at nicks and DSBs, SNV frequencies were normalized to siBRCA2 and siNT2-treated cells, respectively, after subtraction of the background SNV frequency (0.26%) established by analysis of DNA from cells not targeted for cleavage by Cas9D10A. **(B)** Maps of positions and spectra of SNVs within the indicated 65 nt region spanning nicks or DSBs in cells depleted for indicated factors. **(C)** Diagram of mutagenic repair by POLQ of a gap generated by 5’ resection mediated by DNA2/RPA. Dashed gray line, resected DNA.

Mapping identified an asymmetric distribution of SNVs around the cut site at nicks, with most mutations downstream of the nick site; and a symmetric distribution at DSBs. A similar asymmetry was evident in response to depletion of DNA2 or ectopic expression of the dominant negative RPA1-R43E mutant. The distribution of SNVs was not affected by depletion of REV1, REV3, REV7 or POLQ (**Fig 7B**). These results identify an unexpected role for POLQ in generating SNVs at both DSBs and nicks. At nicks, this may occur when POLQ fills in a gap that results from resection by DNA2/RPA, as diagrammed in **Fig 7C**.

## Discussion

Nicks were long discounted as a source of genomic instability, despite their frequency of occurrence. The sequence analysis described here establishes the potential of nicks to promote indels and SNVs, and identifies the mutagenic signatures of DNA2, REV1, REV3, REV7 and POLQ at nicks. These results provide an outline for pathways of repair of DNA nicks mediated by factors familiar from DSB repair which in some cases appear to function differently at nicks than at DSBs.

Nicks are normally protected from mutation by BRCA2, which limits the threat they pose to genomic instability. To permit sequence analysis to be carried out on a dataset that was not limited by a very low frequency of mutations, the pathways of nick repair defined here were largely delineated in cells depleted for BRCA2, which greatly increased the frequencies of indels at nicks as well as frequencies of a-HDR **(Figs. 1, 2**). Availability of an HDR donor protected nicks from deletion. Protection may result from donor annealing that prevents access by DNA2 or other nucleases. The reduction in deletion frequencies caused by provision of a donor exceeded the increase in HDR frequencies, suggesting that donor annealing may protect a nick without leading to productive HDR.

In BRCA2-depleted cells, both deletions and SNVs at nicks were distributed asymmetrically, extending approximately 10 bp 5’ and 20 bp 3’ of the nick site, but no corresponding asymmetry was evident at DSBs **(Figs. 1, 2**). Some of this asymmetry in mutagenesis appears to reflect 5’ resection by DNA2/RPA, which promote mutagenic repair as well as HDR at nicks. DNA2 and RPA are activated in S phase, and analysis of cell cycle dependence of repair suggested that nicks generated in G1 phase may persist until S/G2 phase to undergo resection by these factors (**Fig 3**).

The REV1 dCMP transferase activity appears responsible for most insertions, as their frequency was greatly reduced in response to depletion of REV1, and the great majority were insertions of a single C opposite (**Fig 5**), the signature of REV1 activity. REV1 inserts C on the strand opposite the nick (**Fig. 5C**), which would require the two sides of the nick to be in juxtaposition during this step of repair synthesis. DNA2 not only promoted resection at nicks, it limited 1 bp insertions, which increased 6-fold in response to DNA2 depletion (**Fig 4**). This could occur if resection at the nick by DNA2 generates a gap that becomes the source of deletions, rather than a substrate for REV1-mediated insertion.

REV3 also protects against 1 bp insertions, which increased 3-fold in frequency upon REV3 depletion. This may reflect regulation of REV1 by REV3. REV1 and REV3 are both components of the complex of TLS factors that accompany PCNA during replication. REV1 performs a scaffolding function within this complex by recruiting other polymerases to continue extension that it has begun [25, 41]. Structural studies have documented close and dynamic interactions consistent with the possibility that REV3 may displace REV1 [42], which could prevent 1 bp insertions.

Sequence analysis showed that both REV7 and POLQ limit deletions at nicks (**Fig 6**). REV7 is the non-catalytic subunit of the TLS polymerase POLζ (REV3/REV7), but depletion of REV3 did not affect the deletion frequency, making it unlikely that repair synthesis by POLζ limits deletions. The role of REV7 may instead be architectural, protecting nicks to enable their repair by other factors, as it does at DSBs. POLQ may limit deletion by extending DNA 3’ ends at nicks, as it does at DSBs [35, 43]. In doing so, it may cause SNVs, which are largely POLQ-dependent at both nicks and DSBs (**Fig 7**). POLQ is highly error-prone, generating single base changes at a rate 10-to 100-fold higher than other family A polymerases on undamaged DNA templates in vitro [44]. POLQ is also mutagenic in vivo. It is one of several polymerases that contribute to immunoglobulin gene somatic hypermutation [45, 46] where SNVs arise in the course of mutagenic repair of nicks caused by targeted deamination by AID [1]; and POLQ has also been shown to contribute to the mutation burden in solid tumors [47].

Essentially all SNVs depend upon POLQ (**Fig 7C**). SNVs are distributed asymmetrically around the nick site, extending only 15 bp 5’ but 30 bp 3’ of the nick site. SNVs 5’ of the nick site may be created when POLQ fills in gaps resected by the 5’ nucleolytic activity of DNA2/RPA. SNVs 3’ of the nick site may be created by a similar mechanism, if a factor with 3’ exonucleolytic activity resects nicks to create gaps that can be filled in by POLQ.

The results presented here demonstrate that nicks have considerable potential to contribute to mutagenesis. The increased frequencies of mutagenesis at nicks evident upon depletion of BRCA2 suggests that nick-initiated mutations will be especially frequent in tumors characterized by genetic or regulatory deficiencies that prevent RAD51 loading onto DNA (“BRCAness” [48]). About half of all solid tumors are currently treated with IR, including many BRCA-deficient tumors. IR generates far more nicks than DSBs, suggesting that the mutagenic signatures of these pathways outlined here will be enhanced in tumors treated with radiation.

## Acknowledgments

We thank Brendan Kohrn for assistance with data analysis. This research was supported by R01 CA183967, R21 CA190675, and PO1 CA077852.

## Materials and methods

### Culture and transfection of U2OS cells

U2OS cells were cultured at 37°C, 5% CO_2_ in DMEM (Gibco) containing 10% FBS, 2 mM L-glutamine, 100 units/ml penicillin and 100 μg/ml streptomycin (Gibco # 15140-122). For each siRNA transfection, 3×10^5^ U2OS cells were reverse transfected with 5 pmol of each siRNA (QIAGEN) using lipofectamine RNAiMAX (Thermo Fisher) in 12-well tissue culture plate. At 24 hr post-transfection, cells were trypsinized, washed once with PBS, resuspended in 100 μl of Ingenio electroporation solution (Mirusbio), mixed with 10 pmol premade complex of CRISPR guide RNA and Cas9 (DSBs) or Cas9D10A (nicks) recombinant protein (IDT), transferred to a cuvette and transfected with a 4D-Nucleofector (Lonza), using program CM-104. CRISPR gRNA4 targeted exon 1 of the CD44 gene at the sequence 5’-cctCG<u>TG</u>GCCGCTGAGCCTGGCAC-3’ (PAM sequence is cct, shown in lowercase; cleavage targets the phosphodiester backbone between the underlined <u>TG</u> bases). Parallel transfection with an eGFP expression plasmid (240 ng; Invitrogen) was used to determine nucleofection frequencies (>90%). Following transfection, cells were transferred to 6-well plates and further cultured for 3 days, at which time genomic DNA was isolated with the DNeasy Blood and Tissue Kit (QIAGEN).

For siRNA depletion, 3×10^5^ U2OS cells were incubated with 5 pmol siRNA using lipofectamine RNAiMAX (Thermo Fisher) in a 12-well tissue culture plate for 2hr. Cells were the trypsinized, washed once with PBS, and resuspended in 100 μl of Ingenio electroporation solution (Mirusbio). After addition of 10 pmol premade complex of CRISPR guide RNA and Cas9 or Cas9D10A recombinant protein (IDT) to each well, contents were transferred to a Lonza cuvette and transfection was carried out with the 4D-Nucleofector (Lonza), using program CM-104. Cells were then transferred to a 6-well plate for further culture and harvested at 72 hr post-transfection. Transfection frequencies were determined by parallel transfection with 240 ng of a control eGFP expression plasmid (IDT). Cells were harvested and genomic DNA was isolated with the DNeasy Blood and Tissue Kit (QIAGEN) at 72 hr post-transfection. Parallel analysis of DSBs targeted to the same site provided a physiological control for depletion.

### siRNAs

siRNAs used were siNT2 (4390847), siBRCA2 (s2085), siEXO1 (s17502, S17503) and siMRE11 (s8959, s8960) from Thermo Fisher Scientific; and siDNA2 (SI05067622; SI00370475; SI04293989; SI04309977), siPOLQ (SI02665215, SI00090083, SI00090076, SI0009006); siREV1 (REV1L; SI03090171, SI00115311, SI00115304, SI00115297); siREV3 (SI00045647, SI00045626, SI03058643, SI03120278) and siREV7 (MAD2L2; SI00087710, SI00087703, SI00087696, SI00087689) from Qiagen.

### Amplicon sequencing

Sequencing libraries were prepared followed a modified version of Safe-seq [49] in which genomic DNA underwent limited amplification to add unique molecular identifiers (UMIs), and was then further amplified to generate material for NGS analysis on an Illumina platform.

Primers containing UMIs (5’-TCGTCGGCAGCGTCAGATGTGTATAAGAGACA-GNNNNNNNNNNACTTCGGTCCGCCATCCTCGTC-3’ and 5’-GTCTCGTGGGCTCGGAGAT-GTGTATAAGAGACAGNNNNNNNNNNGCAAATCCCAGCCCTGCTTTC-3’) were used to amplify a 323 bp region of exon 1 of the CD44 gene spanning the cut site for 4 cycles using Phusion Hot Start II DNA Polymerase (Thermo Fisher Scientific). PCR products were purified with Ampure XP beads (Beckman Coulter) and further amplified using a pair of primers containing Illumina Nextera P5 and P7 adapter sites for a total of 25 cycles. Primers for both amplification steps were obtained from IDT. PCR products (480 bp) were purified with 0.7 volume of Ampure XP beads (Beckman Coulter), and reversely cleaned up by 7% Polyethylene Glycol 8000 (PEG) in 875mM NaCl solution (final concentration) followed by a 0.7 volume of Ampure XP beads purification. Purified PCR products were then pooled in equal molar ratio, and sequenced on Illumina MiSeq (600 cycles V3, paired ended reads) at the Fred Hutchinson Cancer Research Center.

Reads were first converted to unaligned SAM files using FastqToSam command from PICARD tools. Then the UMI sequences were extracted and converted to “ab/ba” format where ‘a’ and ‘b’ are the tag sequences from Read 1 and Read 2 using an inhouse script. Remaining reads were then assembled to BAM files in which reads having the same UMI were assembled to consensus sequences using single strand consensus sequence (SSCS) assembly in FASTQ format. Remaining reads were aligned with the reference sequence and analyzed using two established methods. (1) In an approach designed to accurately call SNVs by scoring only mutations found at complementary positions on both DNA strands, SSCS read 1 and read 2 were aligned by Burrows-Wheeler Aligner (BWA) MEM and realigned around INDELs by GATK. Variants were called using Samtools mpileup. INDELs were normalized using bcftools. Output of mutation frequencies, SNVs, insertions, deletions at each position was generated by a python program derived from [50]. (2) SSCS reads were analyzed using CRISPResso2 [51].

INDELs and SNVs from the VCF files were then analyzed using Excel. Sequence analysis of uncut DNA enabled quantification of background frequencies which were subtracted from reported frequencies prior to displaying data using GraphPad Prism. Libraries from several independent experiments were sequenced and analyzed; representative data from a panel of libraries is presented here. Libraries analyzing the effects of depletion of these factors and of DNA2 were constructed concurrently, so frequencies of mutagenesis could be directly compared. Background frequencies for this representative panel of libraries were: insertions, 0.47%; deletions, 0.3%; SNVs, 0.26%. These frequencies were subtracted from results shown.

### HDR reporter assays

Cell culture, transfection, depletion and reporter assays using the Traffic Light (TL) construct in 293T TL cells were carried out as previously described in detail [52]. In brief, the CRISPR RNP was delivered using RNAiMAX according to the manufacture’s recommendation (IDT). To determine nucleofection or RNAiMAX transfection frequencies, Alt-R CRISPR-Cas9 tracrRNA-ATTO 550 (IDT) was used. Transfection frequencies were typically greater than 90%. Unless otherwise specified, DNA DSBs or nicks were targeted by gRNA9 to the sequence 5’-TAAAGCTAAGAGCTCA<u>CC</u>TAcgg-3’ (PAM lowercase, target site between underlined bases). Donors for HDR were either the duplex plasmid pCS14GFP [3] or the 99 nt single-stranded oligonucleotide SSO-2, complementary to the strand nicked by gRNA9. The sequence of SSO-2 is shown below, where uppercase letters denote arms of homology between SSO-2 and the target, and lowercase letters indicate the central region of heterology that must replace sequence in the target to generate a functional GFP gene:

SSO-2: 5’-TGGACGGCGACGTAAACGGCCACAAGTTCAGCGTGTCCGGC-gagggtgagggcgatgcCACCTACGGCAAGCTGACCCTGAAGTTCATCTGCACCACCG-3’

Frequencies of GFP+ cells (HDR) were determined three days post-transfection by flow cytometry, and normalized for transfection efficiency as determined by parallel transfection of a GFP expression plasmid. Data were collected using a BD Biosciences LSR II Flow Cytometer. Each set of assays was performed in triplicate, and a mean frequency of HDR was determined. The values presented represent the mean ± SEM from a representative experiment. Two-tailed T-tests were performed using Microsoft Excel (2015) to determine if the differences between HDR and mutEJ frequencies at different stages of cell cycle were statistically significant. Results are presented as frequencies of HDR among transfected cells, as measured in parallel by transfection with a GFP expression construct.

### Expression constructs

RPA1 expression constructs were generated as derivatives of a RPA1-WT expression construct pLX304-hRPA1 bearing bearing an N-ter V5 tag (Addgene Plasmid #25890). The R41E, R43E, and R41/43E mutations were made by site-directed mutagenesis [53] using the following primers (mutation sites underlined): for hRPA1-R41E: 5’-CCGCCG<u>GAA</u>TATCGACTGCTCATGAGTGATGGATTGAACACTCTATCC-3’ and 5’-AGCAGTCGATA<u>TTC</u>CGGCGGACTATTCCCCGTAGTAATGGGACGGATGTTG-3; for hRPA1-R43E5’-CCGCCGCGTTAT<u>GAA</u>CTGCTCATGAGTGATGGATTGAACACTCTATCC-3’ and for hRPA1-R41/43E, 5’-AGCAG<u>TTC</u>ATAACGCGGCGGACTATTCCCCGTAGTAATGGGACGGATGTTG-3’; 5’-CCGCCG<u>GAA</u>TAT<u>GAA</u>CTGCTCATGAGTGATGGATTGAACACTCTATCC-3’ and 5’-AGCAG<u>TTC</u>ATA<u>TTC</u>CGGCGGACTATTCCCCGTAGTAATGGGACGGATGTTG-3’.

C-terminal CDT1 and GEM tags were added using a modified overlap extension PCR cloning method [54]. The CDT1 and GEM tags were amplified from pCDNA-Cas9-CDT1 using the primers 5’-GCCTATCCCTAACCCTCTCCTCGGTCTCGATTCTACGAGCGGTGGAGGCGGTTCACGCCAA-TTCGCCACCCCCAGCCCCG-3’ and 5’-CTTAACGCGCCACCGGTTAGCGCTAGCTCA-TTACTAGATGGTGTCCTGGTCCTGCGCGGATG-3’ to amplify the CDT1 tag from pCDNA-Cas9-CDT1; and from pCDNA-Cas9-GEM using the primers 5’-GCCTATCCCTAACCCTCTCCTCGGTC-TCGATTCTACGAGCGGTGGAGGCGGTTCACGCCAATTCGCCACCATGAATCCCAGTATGAAGCA GAAAC-3’ and 5’-CTTAACGCGCCACCGGTTAGCGCTAGCTCATTACTACAGCGCC-TTTCTCCGTTTTTCTGCC-3’. The amplified CDT1 and GEM tags were then integrated into pLX304-hRPA1 by PCR. Constructs were verified by both restriction digestion and sequencing; and cell cycle regulation of protein stability was verified by flow cytometry (**S2 Fig**).

To generate the pCDNA-Cas9-CDT1, pCDNA-Cas9-GEM, pCDNA-Cas9D10A-CDT1 and pCDNA-Cas9D10A-GEM expression constructs, we replaced the T2A-BFP tag in both pCDNA-Cas9-T2A-BFP and pcDNA-Cas9D10A-T2A-BFP [3] with the mKO2-hCDT1(30-120) and mAG-hGEM(1-110) cell cycle tags [21]. Cloning was carried out as follows: First, the MfeI site in both pCDNA-Cas9-T2A-BFP and pCDNA-Cas9D10A-T2A-BFP was destroyed by MfeI digestion, fill-in and religation. The plasmids were then digested with NotI and XbaI, to remove the T2A-BFP cassette, which was replaced with a short duplex, Linker MCS, which carries NotI-MfeI-HpaI-NheI sites, and a linker encoding a pentapeptide (Gly-Gly-Gly-Gly-Ser) between the NotI and MfeI sites. We previously used this approach to confer cell cycle restriction to nuclear stability of Activation-Induced Deaminase (AID) [55], and these tags are referred to previously and herein as the CDT1 and GEM tags. To generate CDT1-tagged constructs pCas9D10A-mKO2-CDT1, pCas9-mKO2-CDT1, and Cas9 and Cas9D10A expression plasmids (see above) were digested with MfeI and NheI, and an EcoRI/XbaI fragment bearing the CDT1 cassette from pCSII-EF-mKO2-hCDT1(30-120) [21] was inserted between those sites. To generate GEM-tagged constructs pCas9D10A-mAG-GEM and pCas9-mAG-GEM, the CDT1-tagged constructs were digested with NheI, partially filled in using only dCTP and dTTP, then digested with MfeI to remove the cassette bearing mKO2 and the CDT1 tag; and ligated to a cassette carrying the GEM tag, generated by digestion of pCSII-EF-mAGhGEM(1-110) [21] with HinDIII, partially filled in using only dGTP and dATP, then digested with EcoRI. To generate constructs pCas9D10A-CDT1, pCas9D10A-GEM, pCas9-CDT1 and pCas9-GEM, cassettes encoding the mAG and mKO2 fluorescent proteins were removed by digestion with NotI, overhangs filled to maintain reading frame, and plasmids religated. All constructs were verified by both restriction digestion and sequencing; and cell cycle regulation of protein stability was verified by flow cytometry (**S3 Fig**).

### Cell cycle analysis by flow cytometry

To analyze ectopic expression of CDT1- or GEM-tagged RPA1-R43E, 1×10^6^ U2OS or HEK293T cells were transfected with 750 ng of plasmid, and at 36 hr post-transfection cells were harvested by trypsinization (0.05%), washed twice with cold PBS. Cells were then fixed and permeabilized by incubation with 500 μl of 1x Foxp3 /Transcription Factor Fixation/Permeabilization solution (Invitrogen) for 30 min at room temperature; washed twice with the same buffer; and resuspended in 500 μl of the same buffer. Samples were then divided into two 250 μl aliquots and stained at room temperature for 1 hr with anti-RPA1 antibodies (rabbit, Abcam ab79398; 1:100) to detect endogenous RPA1 or anti-V5 tag antibodies (mouse, BioRad; 1:100) to detect ectopically expressed RPA1. Cells were then washed once in 1 ml PBS and stained with DAPI (10 μg/ml, ThermoFisher) in 10 mM Tris, pH 7.4, 150 mM NaCl, 2 mM CaCl_2_, 22 mM MgCl_2_, 0.1% NP-40, 0.05 mg/ml BSA and 10% DMSO, and then resuspended in 200 μl PBS containing 1% FBS and analyzed by flow cytometry on a BD LSR II instrument, set to record 50,000 cells per sample. Data were analyzed using FlowJo software (version 9.6).

To analyze cell cycle dependence of expression of CDT1- or GEM-tagged Cas9D10A, 3×10^5^ U2OS or HEK293T cells were transfected with 200 ng of expression plasmid, and at 36 hr post-transfection cells were harvested by trypsinization (0.05%), washed twice with cold PBS, then stained with DAPI and further analyzed as above.

## Supporting information: figure legends

**S1 Fig. Treatment with siDNA2 inhibits both a-HDR and c-HDR.**

**(A)** Effects of DNA2, EXO1 or MRE11 depletion on frequencies of a-HDR or c-HDR at nicks or DSBs, respectively. Frequencies were normalized relative to frequencies in 293T TL cells treated with siBRCA2 (nicks) or siNT2 (DSBs). Cleavage was targeted by gRNA9 (see panel B) and supported by a cN ssDNA donor for a-HDR (nicks) or a plasmid donor for c-HDR (DSBs).

**(B)** Left, diagram of the cleavage sites for gRNA 2 and gRNA 9 in the TL construct (arrowheads), with the promoter (P) upstream. Right, effects of depletion of DNA on frequencies of HDR at nicks or DSBs targeted to the TL reporter construct in 293T TL cells by the indicated gRNA. HDR was supported at nicks by a donor complementary to the nicked or intact strand (cN or cI, respectively), in cells treated with siNT2, siBRCA2 or siDNA2+siBRCA2, as indicated; or at DSBs by a dsDNA donor, in cells treated with siNT2 or siDNA2. Frequency values represent the mean ± SEM from a representative experiment; and * and *** indicate p<0.05 and p<0.001, respectively, for the frequency difference between indicated sample and sample treated with siBRCA2 (nicks) or siNT2 (DSBs).

**(C)** Working model for the role of DNA2 resection in a-HDR at nicks supported by a cN or cI donor. Results in panel B show that DNA2 promotes a-HDR by both pathways, and the first step shown in resection 3’ of the nick. The cN or cI donors anneal as shown, and processing then generates a heteroduplex which is resolved by replication.

**S2 Fig. RPA1-R43E inhibits replication but not HDR.**

**(A)** Cell cycle profiles of U2OS control cells (siNT2) or cells treated with siDNA2 or siRPA1.

**(B)** Cell cycle profiles of U2OS control cells (mock) or cells expressing RPA1-WT, RPA1-R41E, RPA1-R43E or the RPA1-R41/R43E double mutant.

**S3 Fig. CDT1 and GEM tags restrict protein expression to G1 or S phase.**

**(A)** Cell cycle profiles of 293T cells transfected with Cas9D10A-mKO2-CDT1 or Cas9D10A-mAG-GEM expression constructs, showing either the entire population or the population gated for PE+ (mKO2) or GFP+ (mAG) cells.

**(B)** Cell cycle profiles of cells mock-transfected or transfected with CDT1- or GEM-tagged derivatives of RPA1-R43E, bearing a C-terminal V5 tag, and stained with anti-RPA antibody, which detects endogenous and ectopically expressed RPA; or with anti-V5 antibody, which detects ectopically expressed RPA1-R43E.

**S4 Fig. Single base insertions in the 4 bp region spanning the nick site.**

Above, sequence of predominant 1 bp insertion. Nick site in the U2OS CD44 gene is underlined, insertion shown in red. Below, tables show nucleotides identified at indicated positions in products of repair containing a 1 bp insertion within the 4 bp region spanning nick site, in U2OS cells treated as indicated Positions are numbered −2, −1, +1, +2, relative to the nick site in the reference sequence GT/GC, where the slash marks the site at which the nick targeted to the CD44 gene by gRNA4 cleaves the phosphodiester backbone.

